# Ultraflexible Ion-Specific Nanoelectrodes for Long-Term Potassium Monitoring in Plants

**DOI:** 10.64898/2026.01.30.702757

**Authors:** Zhongli Yang, Mingyu Shu, Haoran Ma, Guanyi Zhu, Hao Sun, Mingliang Xu, Xiaoling Wei, Wei-Mong Tsang, Daiying Chao, Ruxin Li, Fei He

## Abstract

High-precision *in vivo* monitoring of ion fluxes is essential yet challenging studying plant electrophysiology such as growth regulation, signal transduction and stress responses. Existing methods for probing ion dynamics are limited by low sensitivity, high invasiveness that interferes physiological processes, and the inability to accurately resolve ion homeostasis with required spatial and temporal resolution. Here, we introduce ultraflexible, plant implantable nanoelectrode (PINE) arrays manufactured on 1.2-μm-thick polymer substrates, which enable ultrasensitive and selective measurement of ionic current for month-long via scalable nanofabrication techniques. The fabricated PINE arrays have a smaller dimension than typical plant cells as well as less stiffness, facilitating minimally invasive integration with living plant cells. This subcellular-scale plant-electronic interface allows for reliable, selective detection of K^+^ flux with a detection limit of ∼10⁻⁸ M, and thus allows continuous, stable monitoring of tomato stem cells over six weeks, capturing dynamic potassium fluctuations during all key growth stages. More importantly, the method permits long-term, real-time tracking of ion-specific dynamics without disrupting plant cellular structure or altering endogenous ion concentrations. Therefore, PINE provides unprecedented access to ion homeostasis and signaling networks, making it an excellent platform for precision agriculture and a foundational tool for future digital plant engineering.

## 1. Introduction

Ions are basic, essential components of the complex network of biological processes that support plant life, serving not only as structural elements but also as important signaling molecules, and thus their dynamic roles in structure, enzymatic function, and signaling are all critical for plant survival under fluctuating environmental conditions [1–7]. Maintaining ion homeostasis is vital for cellular functions such as enzyme activity, membrane potential, and metabolic pathways, while dysregulation of ion balance causes oxidative stress, impaired growth, and reduced resilience to environmental challenges including drought, salinity, and pathogen attack [2, 5, 8, 9]. This intricate interplay of ions forms a nexus of molecular and physiological adaptations with major implications for agriculture, stress resilience, and ecological balance. Specifically, K^+^ are pivotal for maintaining cellular turgor, stomatal regulation, enzyme activation, pH homeostasis and osmotic gradients that drive water and nutrient transport [10]. Ca²⁺ acts as a universal secondary messenger, converting environmental signals into biochemical responses [11], while understanding Na⁺ distribution during salinity stress may help improve crop resilience in saline soils [12]. Moreover, Mg²⁺ is the central metal ion in chlorophyll and is essential for light absorption and photosynthetic electron transport [13]. Thus, the balance of ions in plants is a finely tuned orchestra of growth, stress adaptation, and ecological interaction.

Decoding ion mechanisms, from transporter kinetics to microbial interactions, is vital for sustainable agriculture in an era of climate uncertainty[14]. However, real-time measurements of ion dynamics in living plant at the sub-cellular scales remain currently unmet. Traditional methods, such as bulk biochemical assays and static imaging, lack the spatiotemporal resolution needed to directly observe dynamic interactions in living cells [3, 15, 16]. For instance, optical imaging using ion-sensitive probes or sensors that emit light upon binding to specific ions offers a powerful approach for non-invasive ion tracking at the subcellular levels from mm to ms scales [17–23]. Nevertheless, phototoxicity and photobleaching limit long-term imaging, and probe specificity and sensitivity differ depending on the ion type. In contrast, isotope labeling tracks metabolic pathways and redistribution of plant metabolites via mass spectrometry, but it requires specialized instrumentation and expertise, and its temporal resolution is inherently limited for fast dynamics. Electrophysiological techniques measure ion currents via microelectrodes or voltage clamps to study millisecond-scale signaling and decode ion channel dynamics. For instance, patch clamp, ion-selective microelectrodes and organic electrochemical transistors (OECT) directly quantify pH, ion fluxes, sucrose, CO_2_ and reactive oxygen species in leaves, stems, roots or guard cells [24–28]. However, the invasive sample preparation disrupts natural tissue architecture. While wearable sensors allow for non-invasive monitoring of extracellular ion gradients in intact tissues, their spatial resolution is limited for bulk tissue measurements [6, 29–35].

While flexible, implantable electronic interfaces have advanced significantly for long-term, immune-compatible monitoring in animals [36–42], similar progress in plant bioelectronics remains unexplored. Plants present unique challenges—including rigid cell walls, complex vascular systems, and distinct electrophysiological environments—making the long-term stability, biocompatibility, and functional integration of nanoelectronic devices in living plant tissues still unassessed. Nowadays, there is growing demand for high-resolution, real-time ion flux monitoring tools at sub-cellular levels [6, 15, 16, 22, 43, 44], to understand how specific ions regulate fine-scale biological functions under varying environmental conditions and to advance plant electrophysiology. In the current study, we developed ultrathin plant implantable nanoelectrode (PINE) arrays that minimally disrupt the plant cellular structure and function while maintaining high sensitivity and selectivity for monitoring potassium ions. The electrodes are directly integrated with living plant, facilitating unparalleled real-time monitoring of K^+^ flux, which are crucial for osmoregulation, enzyme activation, and membrane potential maintenance. The prolonged lifecycle observation period produced clear and reliable data which validated both the stability and accuracy of the measurements.

## 2. Results

### 2.1. Engineering PINEs for minimally destructive ion-specific biosensing

The schematic illustration presents an *in-situ*, time-resolved continuous monitoring approach for detecting ion flux variations in live plants in response to stressors, as shown in **Figure 1A**. This measurement is performed using a three-electrode system based on potentiometric electrochemical principles, incorporating our custom-designed PINE device as minimally destructive and implantable biosensing probes. The PINE probe is engineered to be sufficiently small (*e.g.*, 50 μm × 1 μm in cross-section) to allow implantation into plant stem tissuesof tomato plants. The sensing site of the PINE probe consists of a gold contact, an electroplated PEDOT-PSS-CNT layer, and an ion-selective membrane (ISM), as detailed in the inset of **Figure 1A**. The ISM is pre-coated via drop-casting to ensure selective ion permeation (**Materials and Methods**). Since the PINE probe directly contacts the plant tissue, ion-selective transport produces a diffusion-limited current that is directly proportional to ion flux, and at the solid-state contact layer between the ISM and the electrode, the ionic signal is converted into electrons, which are then collected by an electrochemical workstation and displayed in real time. Therefore, the system is ideally suited for real-time, high spatiotemporal resolution analysis of ion dynamics under abiotic and biotic stress.

**Figure 1.**
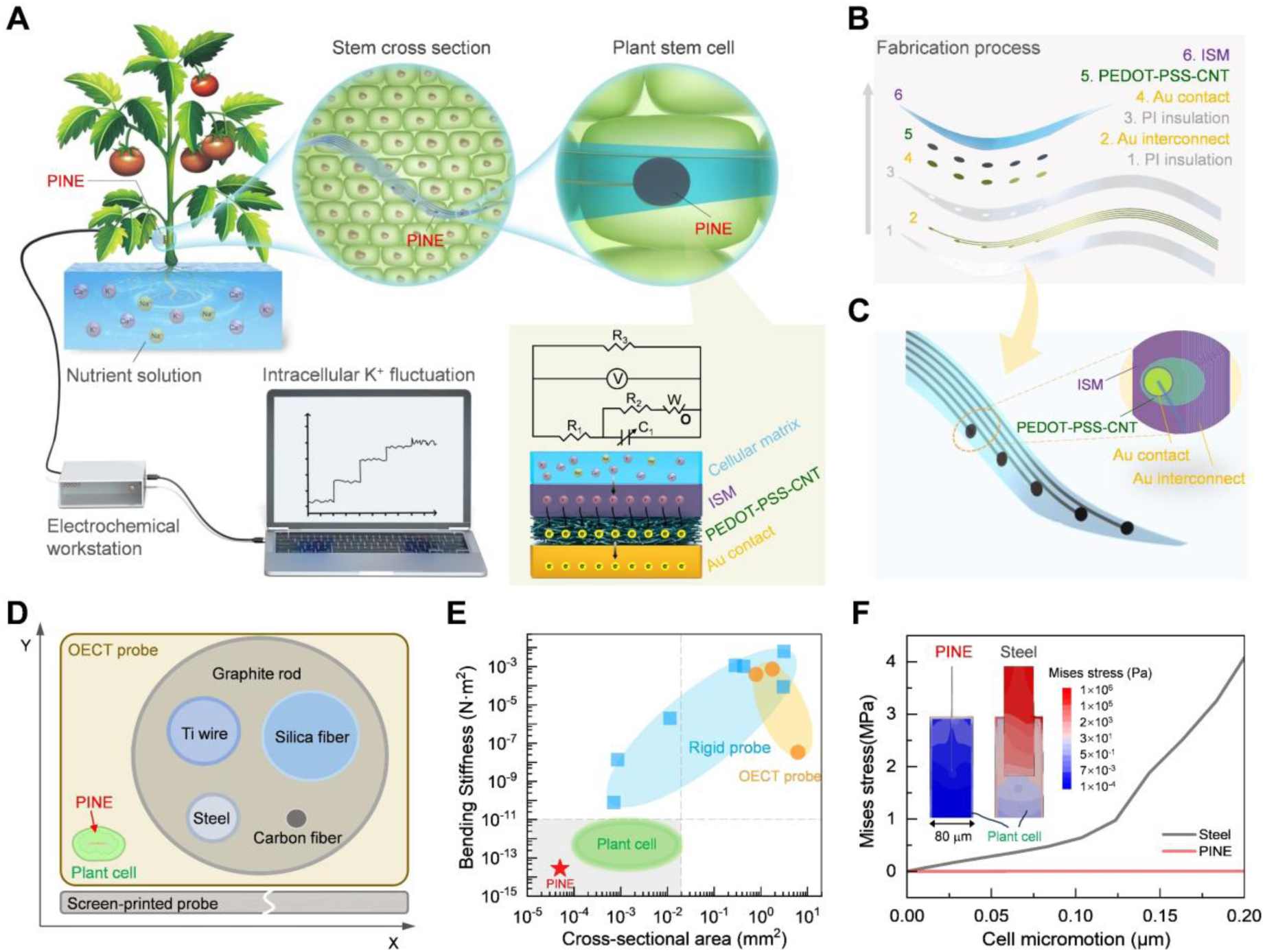
Principle and design of plant implantable nanoelectrodes (PINE). **(A)** Schematic showing PINE implanted into the tomato stem for detecting K^+^ signals. The inset illustrates the mechanism of the PINE/plant interface and presents the equivalent circuit diagram for the ion flux detection. **(B)** Schematic illustration of the lithographic fabrication process of PINE. **(C)** Schematic illustration of thin-film flexible PINE arrays. Inset: magnified view of a single PINE site. **(D)** Schematic illustration comparing the cross-sectional dimensions of a single plant cell, PINE, and previously reported plant probes used for in vivo K^+^ signal measurement. **(E)** Comparison of the implanted cross-sectional area and bending stiffness among previously reported state-of-the-art rigid plant probes (square), OECT plant probes (circle), plant cells (ellipse), and PINE (star) in **D**. In the calculation of bending stiffness, all probes are assigned a uniform length of 1 mm. The detailed parameters of probe materials and dimensions in **D** and **E** are provided in **Table S1**. **(F)** Mechanical analysis comparing the displacement of PINE and stainless steel under 0.2 µm lateral micromotion of a single plant cell. The inset displays the Mises stress profiles within plant cells for pine and steel, respectively, under a lateral micromotion of 0.2 μm in a single plant cell.

To minimize damage to plant tissues and cells while ensuring full electrode insertion, the PINE probe’s dimension is designed with a thickness of ∼1 μm and a width of 50 μm, both of which are much smaller than the average tomato stem cell size (∼100 μm × 200 μm × 200 μm). Moreover, each PINE electrode has an elliptical contact with a short axis of 30 μm and a long axis of 50 μm, designed to maximize ion detection sensitivity. the design of the PINE probe includes 32 individually addressed PINE contacts distributed over three shanks: 10 on one shank and 11 on each of the other two, with a shank spacing of 400 μm. More importantly, this PINE probe was deliberately engineered for plant cell sensing and is distinct from our previous designs[45, 46]. **Figure 1B** illustrates the fabrication process of PINE: the bottom and top insulation layers, each 500 nm thick, together with the Au interconnects and contacts (steps 1 – 4), are first fabricated by standard 4-inch MEMS lithography, after which a custom printed circuit board (PCB) is mounted on the appropriate Au contact pads. The implantable segment is then immersed in a nickel etchant for 5–10 minutes to release the flexible portion. To improve its electrochemical performance, in particular the conversion efficiency of ion signals, the electrode surface is electroplated with a composite of carbon nanotubes (CNTs) and poly(3,4-ethylenedioxythiophene): poly(sodium 4-styrenesulfonate) (PEDOT-PSS) (step 5). Finally, the PINE structure is formed by drop-casting the modified electrodes with a prepared K^+^ ISM mixture solution (step 6) (see **Figure 1C, Materials and Methods**).

We compare the geometrical and mechanical properties of our designed PINE probe with those of previously reported state-of-the-art implantable electrodes, including organic electrochemical transistors (OECTs) and rigid electrodes such as stainless steel, titanium wire, graphite rods, carbon fiber, and silicon fiber, for the detection of various chemicals, such as ions, molecules, and radicals in living plants, as shown in **Figure 1D, E**. We define the implantation cross-sectional dimension as the two-dimensional geometric measurement obtained when a perpendicular cut is made through the electrode at the specific location where it interfaces with the plant tissue. It should be noted that our PINE probe exhibits the smallest implantable size among all reported implantable plant sensors, even one order of magnitude smaller than the average size of plant cells. While OECTs are highly sensitive in detecting ions and metabolites at physiological concentrations, their relatively large size compared to nanoscale sensors limit their applicability in confined biological spices. Furthermore, certain OECT materials degrade when exposed to moisture and biological fluids over time, which may compromise their long-term reliability. The bending stiffness values were calculated according to the formulas (**Equations S1, S2**). As shown in **Figure 1E**, compared with rigid electrodes and flexible OECTs, PINE also exhibits the lowest bending stiffness, especially as low as 2.8 × 10⁻¹⁴ N·m², and smallest cross-sectional area. Since both the stiffness and dimensions of a single plant cell are slightly larger than those of PINE, it follows directly that implanting PINE into plant tissue is not only feasible but also highly advantageous, because its small size and great flexibility make it ideally suited to fit within the cellular environment without perturbing normal cell functions.

To further investigate the mechanical interactions between the PINE probe and plant cells after implantation, we developed a finite element analysis (FEA) model to simulate and compare the stress associated with micromotion for both the PINE probe and a 30-μm-diameter steel wire inserted into a plant cell (**Materials and Methods**, and **Table S2**), as illustrated in **Figure 1F**. In the simulation, a single plant cell was approximated as having dimensions of ∼80 μm × 150 μm and a cell wall thickness of ∼5 μm. The Young’s moduli of the cellular matrix and cell wall were selected accordingly to be 1 kPa and 100 MPa respectively, in general, reflecting the mechanical properties of the cellular contents and the more rigid outer cell wall structure. The simulation results shown that even during a micromotion of 0.2 μm, the stress and strain induced by the PINE probe within the plant tissue are 4–6 orders of magnitude lower than those caused by metal wires. Again, this striking disparity underscores the PINE probe’s superior mechanical compatibility with the delicate plant cell structures.

The drastic reduction in mechanical stress is clearly and logically explained by the fact that the PINE probe has both an excellent structural design and an appropriately chosen material composition, allowing it to mimic the flexibility and elasticity of biological tissue, therefore minimizing mechanical mismatch that could lead to tissue deformation or damage. As a result, the probe interacts gently with the cell, preserving cellular integrity and function. This is particularly advantageous for plant cells, which are inherently stiffer and more structurally complex due to cell walls and large vacuoles.

### 2.2. High-performance ion flux detection using PINE probes

**Figure 2A, B** show the Schematic illustration of the PINE probe fabrication process and wafer-scale PINE probes and a close-up view of individual probe structures, respectively. The overall thickness of the implantable part is measured to be ∼1.2 μm (**Figure S1**). The released implantable component of the PINE device is ultrasmall and provides a comfortable fit for the leaves and stems of tomato plants (**Figure 2C, D**). As shown in **Figure 2E**, an optical micrograph captures a single released shank containing ten PINE contacts suspended in water. The image confirms that the PINE probe bends freely in water, demonstrating its mechanical flexibility. Focused ion beam (FIB) milling combined with scanning electron microscopy (SEM) was employed to accurately assess the structural morphology and thickness of each layer following the PEDOT-PSS-CNT electroplating process, enabling precise observation of the deposited materials and their interfacial bonding. As shown in the cross-sectional SEM image presented in **Figure 2F**, a densely adhered thin layer of PEDOT-PSS-CNT, ∼100-150 nm thick, was successfully formed on the Au contact, acting as a highly efficient ion-to-electron transducer in the PINE. We choose PEDOT-PSS-CNT due to its large specific surface area, which enhances charge storage; low interfacial impedance, which enable efficient signal transmission; and strong electron-ion conversion capability, which bridges ionic signals in live plant with electrical signals in PINE. The layer thickness of 100-150 nm was chosen to optimize performance; it provides sufficient thickness to maintain moderate charge capacity for reliable signal conversion, while remaining thin enough to minimize polarization effects that could distort the signal.

**Figure 2.**
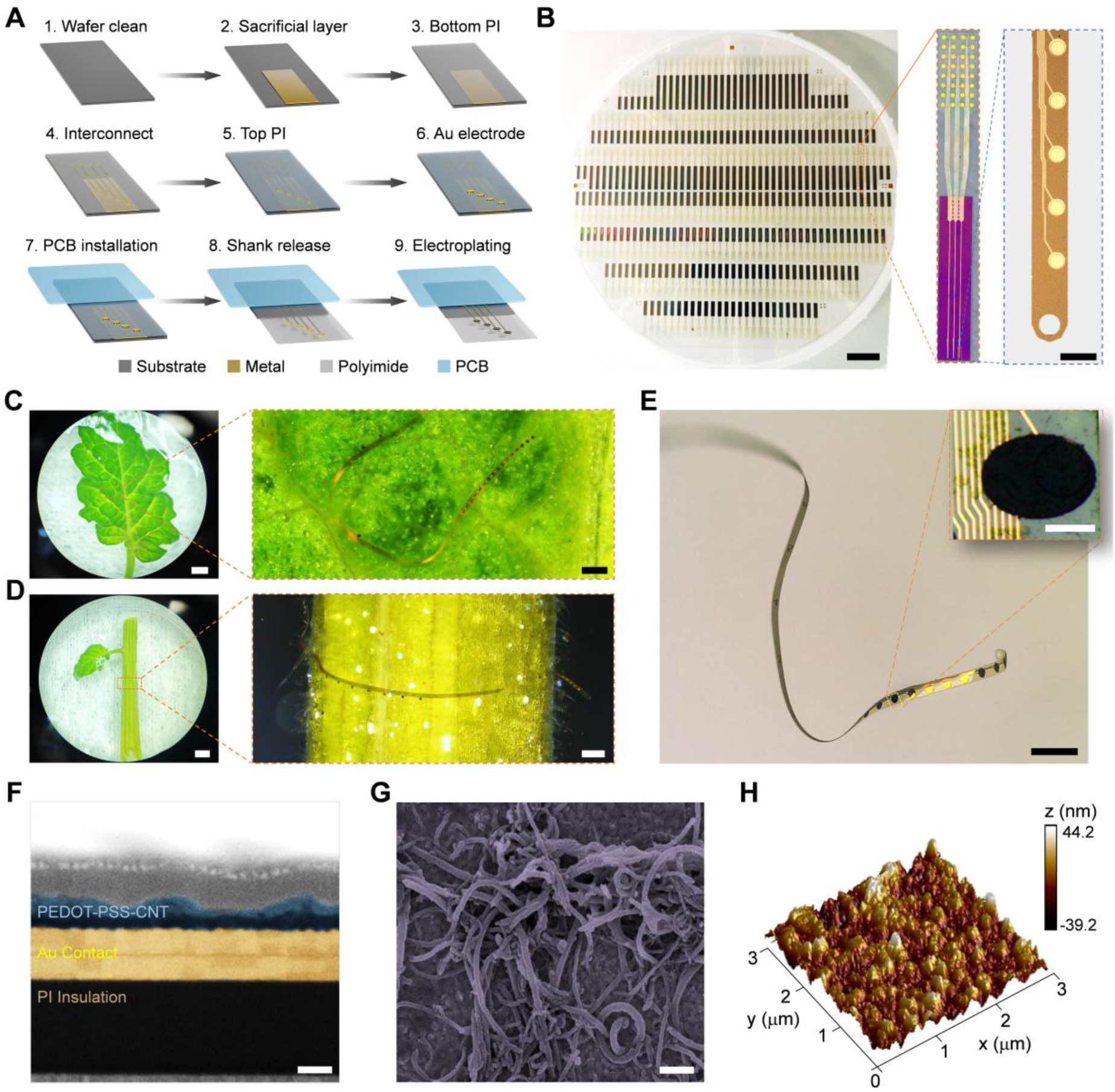
Fabrication and characterization of PINE. **(A)** Schematic illustration of the PINE probe fabrication process. **(B)** Wafer-scale fabrication of PINE plant implantable electrodes performed on a 6-inch fused silica substrate. Each wafer contains 286 probes. Insets show a close-up digital image of the PINE probe retrieved from the wafer substrate, as indicated by the orange box, and an optical micrograph of the implantable part of the PINE probe. Scale bars: 10 mm and 50 μm (inset). **(C, D)** Photographs of the flexible parts of the PINE probe floating on the leaf and wrapping on the stem of a tomato plant, respective. Insets show the optical micrographs as indicated by the orange box, respectively. Scale bars: 2 mm and 200 μm (inset). **(E)** Optical micrograph of a PINE probe floating on water, demonstrating its flexibility. The inset shows an optical micrograph of the top view of a single PINE site. Scale bars, 200 μm and 20 μm (inset), respectively. **(F)** Focused ion beam (FIB) and scanning electron microscope (SEM) image showing the cross-sectional structure of a PINE contact in the inset in **B**. Each layer is pseudo-colored and labelled. Scale bar, 200 nm. **(G)** SEM imaging showing the surface morphology of a PINE contact. Scale bar, 200 nm. **(H)** Atomic force microscopy (AFM) image of the PINE surface.

The surface morphology and roughness of PEDOT-PSS-CNT are fundamental factors determining the electrochemical properties of PINE in ion-to-electron transduction, and because they directly influence ion diffusion, charge accumulation, and transduction efficiency, it is logical and necessary to investigate them using SEM for surface morphology analysis and AFM for nanoscale surface roughness quantification. As can be shown in **Figure 2G**, PEDOT-PSS-CNT has a fibrous nanowire-like morphology resulting from the efficient incorporation of CNTs into the PEDOT-PSS matrix, and the well-dispersed, aligned CNTs form an interconnected conductive network that markedly shortens the electron transport distance, hence greatly improving charge transport efficiency. The surface morphology of the modified sample is examined by AFM in **Figure 2H**, giving an average roughness (r. m. s.) of 11.8 nm, indicating improved topography with the incorporation of hydroxylated multi-walled CNTs. Moreover, the presence of CNTs during electrodeposition further amplifies the specific surface area, providing more active regions for electrochemical reactions, which is vital for improving the overall PINE performance. Since PINE’s solid-state contact layer bonds closely with the ISM, maximizing the interfacial contact area can enhance bilayer capacitance, promoting charge accumulation and storage. This larger interface allows more ions to gather and transfer efficiently between the layers, facilitating faster ion-electron interactions.

Cyclic voltammetry (CV) measurements were conducted in a 1 M KCl electrolyte solution to evaluate the electrochemical performance of pristine, PEDOT-PSS, and PEDOT-PSS-CNT coated electrodes. The tests were carried out within a voltage range of 0 V to 0.5 V at a scan rate of 10 mV/s (**Figure 3A**). The PEDOT-PSS-CNT electrode exhibited a nearly symmetric and rectangular CV profile, indicating ideal capacitive behavior. The capacitive currents suggest a high charge storage capacity and efficient ion transport, thus demonstrating the composite’s ability to reversibly store and release electrical energy. The capacitance of the PEDOT-PSS-CNT was 1.2×10⁵ F/mm² (using **Equation S3**), two orders of magnitude higher than that of the other two electrodes. The effective electroactive surface area of the PEDOT-PSS-CNT layer, derived from CV curves (**Figure 3B**), was ∼860 μm² (using **Equation S4**), significantly larger than that of the other two materials. This finding highlights its superior electrochemical activity, enabling more stable and accurate potential measurements with PINE. Additionally, the electrodeposition time of PEDOT-PSS-CNT was also investigated, and the optimal deposition time was determined to be 100 s (**Figure 3C**). As shown in **Figure 3D**, its capacitance increased with the scan rate, confirming good capacitive behavior even at higher rates. Moreover, the anodic current measured at 0.2 V increase linearly with the scan rate (**Figure 3E**), indicating fast ion and electron transport kinetics, which reduces internal resistance and improve PINE response time.

**Figure 3.**
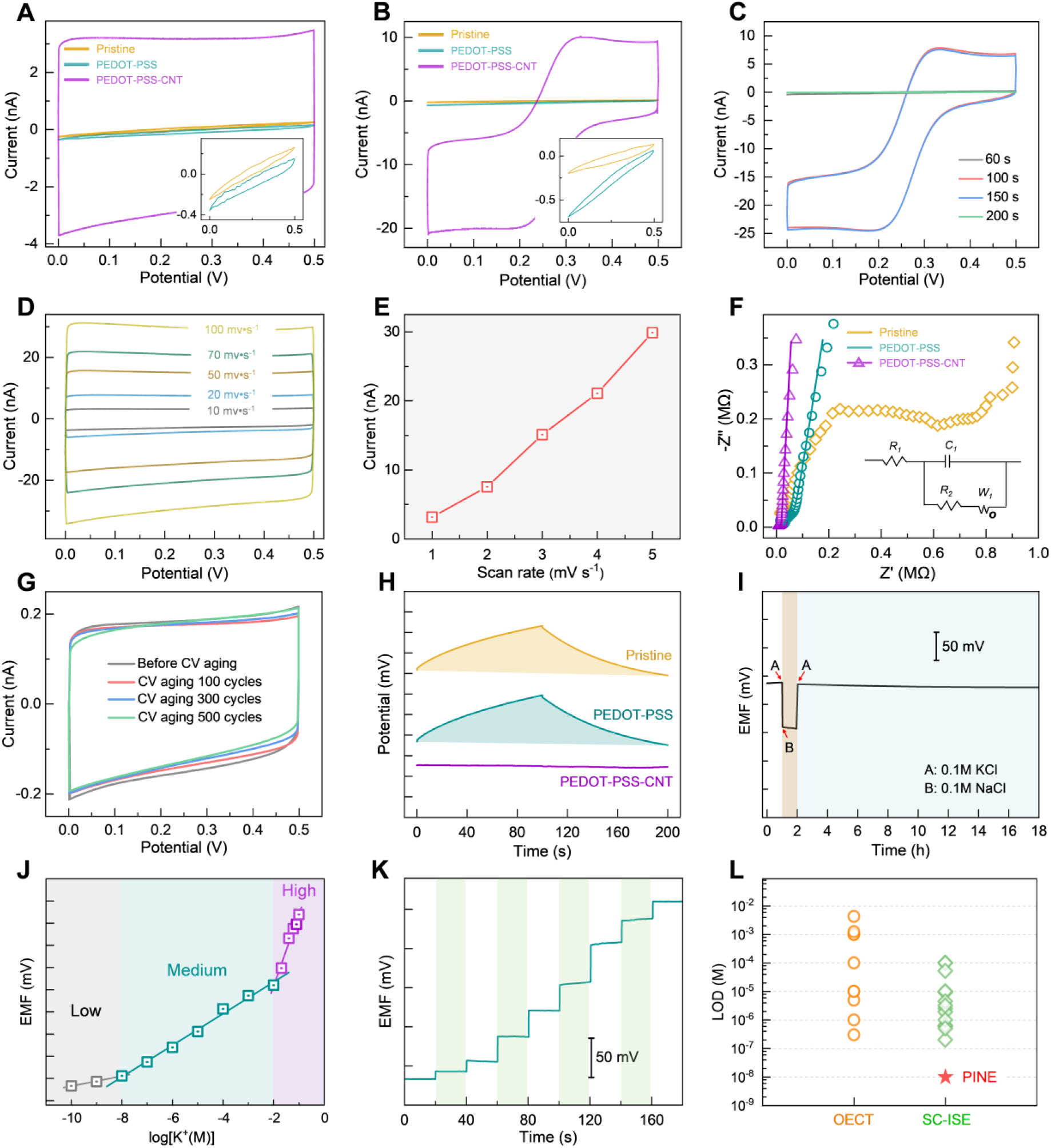
Electrochemical performance evaluation of ion detection using PINE. **(A)** CV curves of the pristine, PEDOT-PSS, and PEDOT-PSS-CNT modified electrodes, respectively. The inset provides an enlarged view of the CV curves for the pristine and PEDOT-PSS modified electrodes. **(B)** CV curves recorded at a scan rate of 10 mV/s in a 1 M KCl solution containing 10 mM [Fe(CN)₆]³⁻ for the pristine, PEDOT-PSS, and PEDOT-PSS-CNT modified electrodes, respectively. The inset presents an enlarged view of the CV curves for the pristine and PEDOTPSS modified electrodes. **(C)** CV performance of the PEDOT-PSS-CNT-modified electrode evaluated at different electrodeposition times (60, 100, 150, and 200 seconds) in 1 M KCl solution containing 10 mM [Fe(CN)₆]³⁻, using a scan rate of 10 mV/s. **(D)** Representative *in vitro* cyclic voltammetry performed at various scan rates (10, 20, 50, 70, and 100 mV/s) demonstrates linearity and stability of PINE. **(E)** Current values observed at different scan rates under a constant voltage of 0.2 V demonstrating the linearity and stability of PINE. **(F)** Nyquist plots of the pristine, PEDOT-PSS, and PEDOT-PSS-CNT modified electrodes, respectively. The inset displays the corresponding equivalent circuit model. **(G)** CV aging curves of PEDOT-PSS-CNT before aging and after 100, 300, and 500 aging cycles, respectively. **(H)** Comparison of chronopotentiograms obtained from the pristine electrode, the PEDOT-PSS-modified electrode, and PINE, measured under an applied current of +1 nA for 100 s followed by −1 nA for another 100 s in 0.1 M KCl solution. **(I)** Water layer tests of the PINE were conducted under 0.1 M KCl and 0.1 M NaCl solutions, demonstrating the excellent performance of the PINE. The red arrows at points A and B indicate the timestamps when KCl and NaCl were applied, respectively. **(J)** Plot showing the calibration of EMF detection by gradually applying KCl solutions ranging from 10^−10^ to 0.1 M. **(K)** Potentiometric response between 10^−10^ and 10^−2^ M KCl. Gray, green, and pink bars indicate the potentiometric response at low (10^−10^–10^−8^ M), medium (10^−8^–10^−2^ M), and high (10^−2^–1 M) KCl concentrations, respectively. **(L)** Comparison of detection limits (LODs) among previously reported state-of-the-art K^+^ ion-selective electrodes (diamond), OECTs (circle), and PINE (star). All corresponding parameters and references are provided in **Table S3.**

The electrochemical impedance spectroscopy (EIS) spectra of pristine, PEDOT-PSS, and PEDOT-PSS-CNT electrodes are shown in **Figure 3F**. The data were fitted using the Randles equivalent circuit model (inset), which incorporate the charge transfer resistance (R_2_) at the electrode-electrolyte interface. The R_2_ value of the PEDOT-PSS-CNT electrode was 3.9 kΩ, much lower than that of the PEDOT-PSS electrode (6.7 kΩ). This reduction indicates faster electron transfer kinetics and enhanced electrochemical performance of PINE. The incorporation of CNTs into PEDOT-PSS matrix improves both electrical conductivity and surface characteristics by establishing efficient conductive pathways that facilitate electron transport and electrochemical reactions — particularly beneficial for applications requiring rapid electron transfer. The stability of the PEDOT-PSS-CNT electrode was evaluated over 500 charge-discharge cycles (**Figure 3G**), revealing minimal current degradation and excellent performance retention. This enhanced stability arises from the synergistic effects of PEDOT-PSS and CNTs, which improve mechanical strength and electrical conductivity.

We employed current-reversal chronopotentiometry to evaluate the short-term potential stability of different electrode coatings. As shown in **Figure 3H**, the typical chronopotentiograms for three different electrode configurations—pristine Au, Au modified with PEDOT-PSS, and PEDOT-PSS-CNT—are presented for comparative analysis. The potential drift, measured as Δ*E*/Δ*t* (change in potential over time), was 2.1 mV/s for pristine Au, indicating poor stability. In contrast, the electrode with PEDOT-PSS-CNT layer exhibited a much smaller drift of 30 μV/s, demonstrating excellent potential stabilization capability. More importantly, the potential stability of the PEDOT-PSS layer without CNTs was essentially the same as that of the uncoated Au electrode (2.2 mV/s), thus unambiguously showing that the CNTs are the key factor in enhancing potential stability. Hence, the conclusion is clear: inserting PEDOT-PSS-CNT between the K^+^-selective membrane and electrode greatly improves the potential stability of PINE.

A water layer between the ISM and solid contact layer severely compromises potentiometric sensor stability during *in vivo* plant measurements. This occurs because hydrophilic solid contact materials trap water at the interface, forming a diffusion barrier that impedes ion transfer. As ionic strength fluctuates over time—especially in long-term or environmentally variable conditions—sensor sensitivity and accuracy degrade. **Figure 3I** shows the PINE electrode’s stable response across three consecutive solutions: 0.1 M KCl → 0.1 M NaCl → 0.1 M KCl. Potential drift is negligible throughout the cycle, enabling sustained operation under long-term maintenance conditions. This stability strongly indicates that the PEDOT-PSS-CNT composite suppresses interfacial water layer formation. The most likely mechanism is enhanced surface hydrophobicity (reducing water accumulation) combined with CNTs’ superior charge transport, jointly improving electrochemical stability.

PINE’s ion flux detection showed no baseline drift or fluctuation over 16 h, confirming its long-term stability, robustness, and reproducibility. Its potentiometric response was also characterized in KCl solutions ranging from 10⁻¹⁰ M to 10⁻¹ M. The results of **Figure 3J, K** and **Figure S2** show that PINE delivers a highly stable and reproducible response across the entire concentration range. Notably, PINE exhibits a near-Nernstian response with an average slope of 35.7 ± 1.1 mV per decade and a strong linear correlation (R² = 0.993) between 10⁻⁸ M and 10⁻² M (**Figure 3K**). The detection limit was determined using the intersection method, which identifies where two linear segments of the calibration curve converge. This approach yielded a detection limit of 10⁻⁸ M, underscoring PINE’s high sensitivity and for trace ion detection. A comparison of PINE’s performance with previously reported potassium ion-selective electrodes (**Figure 3L**) reveals that PINE offers a lower detection limit and broader detection range for K^+^, outperforming both OECT-based and conventional solid-state ion-selective electrodes (SC-ISE). Its low detection limit ensures high sensitivity in low-ion environments, while its wide detection range supports accurate K^+^ measurement in diverse conditions. A detailed comparison of various K^+^-selective electrodes—including sensitivity, detection range, response time, and stability—is shown in **Table S3** in the Supplementary, demonstrating PINE’s competitive advantages.

### 2.3. High-precision long-term monitoring of K^+^ dynamics in tomatoes

PINE was implanted into tomato stems to monitor in vivo potassium concentrations over time. **Figure 4A** shows the experimental timeline across three key growth stages—flowering, fruit set, and fruit maturation—when potassium demand is highest. Given potassium’s critical roles in enzyme activation, osmoregulation, and signal transduction, monitoring began two days post-implantation and continued through all stages to capture real-time K⁺ dynamics. The goal was to assess PINE’s ability to reveal plant nutrient dynamics, with direct relevance to plant physiology and agricultural applications.

**Figure 4.**
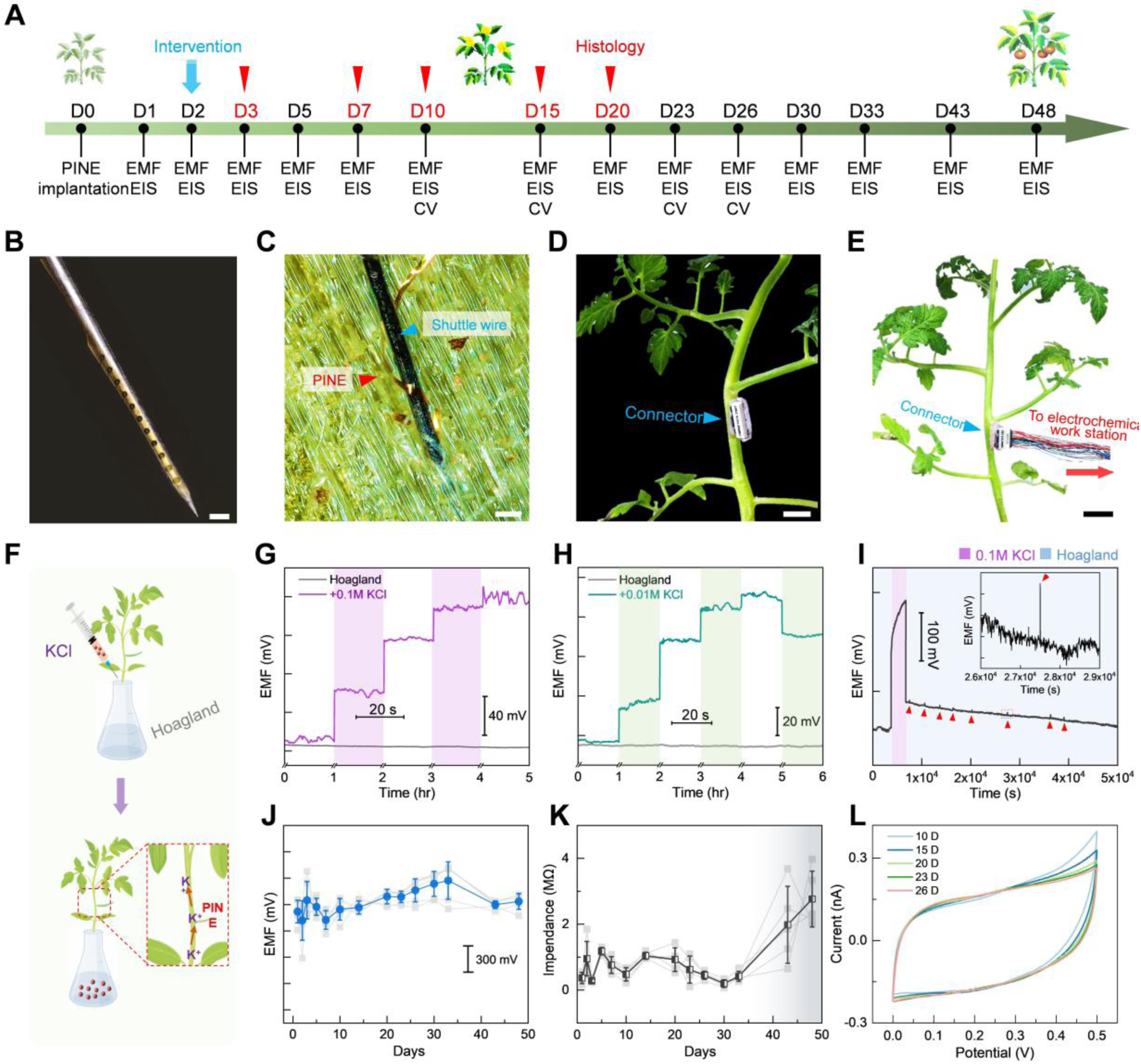
Implantation of PINE into the tomato stem for long-term, high-precision monitoring of potassium dynamics. **(A)** Timeline of *in vivo* electrochemical characterization of PINE and chronic potassium ion monitoring with an implanted PINE probe in tomato stems. **(B)** Optical micrograph of a PINE probe assembled with a tungsten shuttle wire, intended for implantation into the plant. Scale bar, 100 μm. **(C)** Optical micrograph illustrating the implantation of the PINE into the tomato plant stem with assistance from the Tungsten shuttle wire and the subsequent detachment of the PINE from the shuttle in DI water. Scale bar, 100 μm. **(D)** Photograph of a PINE device implanted into the tomato stem, with the backend connector mounted on the stem. **(E)** Photograph showing the measurement of the ion dynamics by connecting the PINE device to the electrochemical work station. **(F)** Schematic illustration of *in vivo* K^+^ monitoring in tomato plants with PINE implanted. Tomato plants with PINE implanted were grown in Hoagland solution, to which a quantitative amount of KCl solution was added using a syringe for each measurement. **(G)** *In vivo* potentiometric response of PINE in Hoagland solution (gray) and a medium concentration of KCl (pink) over time, respectively. **(H)** *In vivo* potentiometric response of PINE in Hoagland’s solution (gray) and a high-concentration KCl solution (green) over time. For the potentiometric measurements in KCl **(G, H)**, the EMF was recorded continuously for 20 seconds immediately after the addition of 0.01 M KCl every 1 hour. **(I)** Transition of the potential observed when tomato plant roots are immersed in 0.1 M KCl and transferred to Hoagland’s solution. Periodic potential bursts are indicated by red triangles. The inset provides an enlarged view of a peak occurring between 2.6 and 2.9 × 10⁴ s. **(J)** *In vivo* electromotive forces in tomato stems with PINE implantation as a function of days post-implantation (*n* = 5 effective sites, mean ± s.d.). **(K)** *In vivo* impedance of the PINE array implanted in the tomato stem, measured at 1kHz, as a function of days post-implantation (*n* = 5 effective sites, mean ± s.d.). **(L)** Representative repeatable CV profiles of a PINE site recorded *in vivo* at different time points post-implantation.

The released section of PINE was temporarily attached to a shuttle device made of tip-sharpened tungsten microwires (50-75 μm in diameter) using a bio-dissolvable adhesive (**Materials and Methods**). This specific diameter range of shuttle wire ensured sufficient stiffness for penetration into the plant stem at various locations while minimizing initial tissue damage. **Figure 4B** shows an optical micrograph of an assembled PINE probe attached to the tungsten wire, ready for implantation into tomato stems. The assembled device was then mounted on a micromanipulator for precise delivery into a fixed tomato stem, and all implantation procedures were conducted under observation using an optical surgical scope. **Figure 4C** shows the PINE probe implanted into a tomato stem with the assistance of a tungsten wire. The insertion depth of PINE is approximately 2 mm, reaching through the epidermis, phloem, xylem, and pith. Since the phloem and xylem are the principal pathways for water and ion transport, their localization makes them ideal sites for sensitive and reliable monitoring of ion dynamics. After insertion, DI water was applied to dissolve the bio adhesive and thereby detach the PINE probe from the tungsten wire, after which the shuttle wire was slowly withdrawn, leaving only the ultra-small, flexible PINE probe embedded in the stem. The insertion footprint was only ∼50 μm in diameter—much smaller than the plant stem cell—causing minimal tissue disruption (**Figure S3, Movie S1**). **Figure 4D-E** shows the PINE device implanted in the tomato stem, with its backend connector attached and ion dynamics measured via connection to an electrochemical workstation. Tomato seedlings with implanted devices had their roots submerged in Hoagland’s solution containing varying KCl concentrations, and corresponding electrical signals were recorded (**Figure 4F**). Potentiometric measurements of K^+^ fluxes in vascular bundles revealed that 0.1 M KCl application induced a gradual 4-h rise in stem K^+^ potential, indicating progressive root uptake and stem translocation. As shown in **Figure 4G**, after 3 h, the potential increase slowed markedly; by 4 h, plants showed wilting and lodging—suggesting rising stem K^+^ coincided with declining water content. Thus, prolonged high-K^+^ exposure harms tomato growth. In contrast to the stable signal in Hoagland’s solution, the PINE system reliably and sensitively detected dynamic K^+^ changes in stems. We repeated the experiment at a lower K^+^ concentration (0.01 M) and observed structural stability—no wilting or lodging—for 4 hours (**Figure 4H**). In the fifth hour, electrical signal measurements revealed a marked drop in potential, indicating that prolonged high K^+^ exposure impairs intracellular function or membrane integrity. Critically, these results confirm that the PINE system reliably detects real-time K^+^ signals in plants, enabling studies of ion dynamics and stress responses.

The stability of PINE’s signal detection in plants was evaluated, as the results shown in **Figure 4I**. First, the plants were immersed in Hogland’s solution for one hour, during which the electrical potential remained stable. Subsequently, the plants were transferred to a mixed solution containing 0.1 M KCl for another hour, which resulted in a notable increase in potential. Despite the excess K^+^ in the external solution, the roots continued to absorb potassium ions, leading to elevated K^+^ levels in the vascular tissues. Over time, the potential continued to rise, suggesting a progressive accumulation of K^+^ from the roots to the stems. When the plants were returned to Hogland’s solution for 12 hours, the potential dropped significantly but remained slightly above the initial level, gradually declining thereafter. This phenomenon may be attributed to the excessive K^+^ intake impairing ion transport channels and slowing the re-translocation of K^+^ from the stems to roots. Interestingly, small potential bursts occurred every 50 minutes, indicating periodic fluctuations in K^+^ concentration. The underlying mechanism of this K^+^ bursting behavior remains to be elucidated. Long-term PINE performance in tomato stems was monitored for over one month (**Figure 4J**). Initial potential fluctuations during the first 10 days likely resulted from implantation-induced tissue damage. Stable readings were achieved from day 26 onward—indicating full wound healing and physiological equilibrium. Minor fluctuations on day 34 aligned with normal plant growth, while the potential decline by day 48 coincided with natural senescence following flowering and fruiting. These results confirm PINE’s long-term stability and reliable functionality in living plants.

**Figure 4K** shows impedance changes at 1 kHz over time: (1) Frequent fluctuations during days 0–10 reflect active wound healing and environmental adaptation; (2) Stable impedance from day 10 to day 33 indicates physiological equilibrium—uniform tissue structure and stable water content; (3) A sharp rise after day 33 signals senescence, driven by cell degradation, declining water content, loss of turgor pressure, and impaired membrane integrity—classic hallmarks of plant aging. PINE’s in vivo electrochemical performance in tomato plants (**Figure 4L**) remained stable over time: the CV curve area at a representative implant site showed no significant change post-implantation. This confirms rapid stabilization and excellent long-term electrochemical stability in plant tissue—enabling reliable, real-time monitoring of biological processes without degradation.

### 2.4. Histological analysis of the PINE/plant interface shows their seamless integration

We investigated the nature of the PINE/plant interface through histological analysis at various time points after implantation. A schematic of a PINE probe inserted into the tomato plant stem (**Figure 5A**) illustrates the orientations of the tissue sectioning, which was performed either parallel or perpendicular to the implanted probe at different times post-implantation. In general, tissue slices were prepared by fixing the plant tissue using standard procedures (**Materials and Methods**), with the PINE probes left in place. In contrast, the control tungsten wire was retracted to facilitate slicing. **Figure 5B** shows an optical micrograph of a PINE probe successfully implanted in a tomato stem, confirming its near-straight configuration post-implantation. **Figures 5C, 5D**, and **Movie S2** show that two weeks after implantation, the probe remains inside tomato stem with normal morphology and intact cell. Thus, PINE retains structural integrity and functionality across diverse biological environments—a key advantage for plant physiology, where tissue composition changes dynamically during development and in response to environmental conditions..

**Figure 5.**
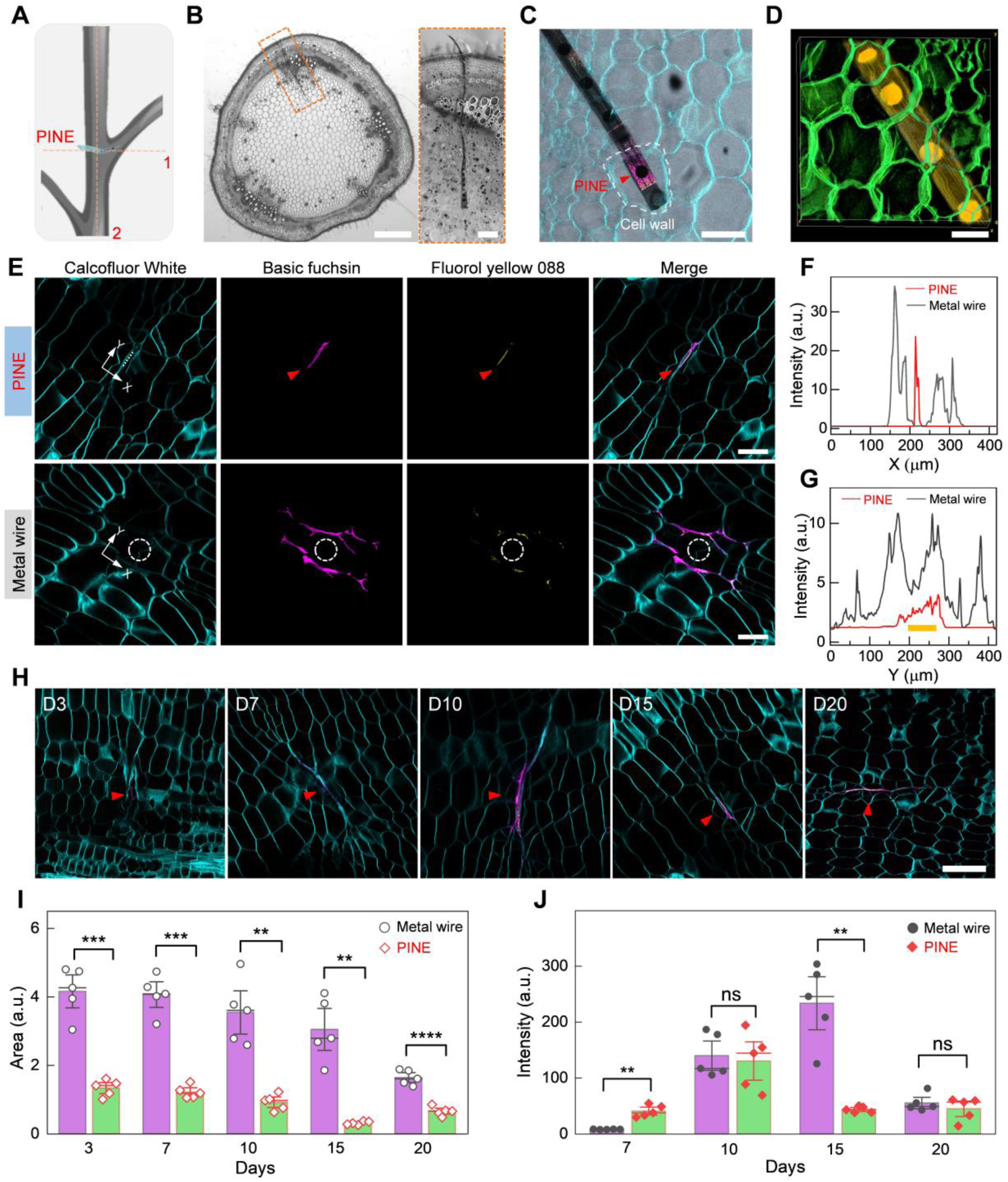
Implanted PINE-plant histology. **(A)** Schematic illustration of the preparation of tomato plant stem slice samples. The orange dashed lines indicate the slicing directions: parallel (line 1) and perpendicular (line 2) to the implanted PINE, respectively. The histology sections in **B**-**D** were sliced along line 1, and those in **E**-**J** were sliced along line 2. **(B)** Digital picture of a tomato stem slice containing a PINE probe. The orange dashed box delineates the site of PINE implantation. Inset provides a closed-up view of the optical micrograph. Scale bars: 1 mm, 200 μm (inset). **(C)** Representative confocal image taken at a depth of 50 μm in a 300-μm thick stained stem slice shows a stem slice containing multiple PINE sites. The white dashed line outlines a cell wall, and the red triangle indicates a PINE site inside the plant cell. See **Movie S2** for a full view of **C**. Scale bar: 100 μm. **(D)** 3D reconstruction of a 75-μm thick stained slice by confocal imaging shows PINE integrated into tomato stem. Scale bar: 50 μm. **(E)** Confocal micrographs showing suberin and lignin staining in cross-sectional slices of a PINE probe (top) and the surrounding area of a 75-µm-diameter steel (bottom), both implanted in the tomato plant stem 15 days post-implantation. The red triangles indicate the cross-sections of the PINE contained in the slice, and the white dashed circles outline the implantation sites of a 75-µm-diameter steel. Scale bar, 100 μm. **(F, G)** Normalized fuchsin fluorescence (purple) intensity profiles were plotted along the *X* (**F**) and *Y* (**G**) axes of the confocal images in **E**. The orange bar indicates the position of the PINE cross-section. **(H)** Time-dependent confocal images of suberin and lignin-stained PINE-plant interfaces at 3 days, 7 days, 10 days, 15 days and 20 days post-implantation. The red triangles indicate the cross-sections of the PINE contained in the slice. Scale bar, 100 μm. **(I)** The normalized area of fuchsin fluorescence (purple) around the site of a 75-μm diameter metal wire and the PINE probe at different period post-implantation at different period post-implantation. **(J)** Comparison of the normalized fuchsin fluorescence intensities (purple) around the site of a 75-μm diameter metal wire and the PINE probe at different period post-implantation. All error bars represent ±1 s.e.m. In **I, J**, each condition was independently repeated across *n* = 5 distinct tissue volumes implanted with metal or PINE, respectively. Additional data are shown in **Figures S4-S7**. All tissue slices were prepared post-implantation into the stems of tomato plants (see **Materials and Methods**). Significance levels are indicated as follows: n.s., no significance; \*\**P* < 0.01; \*\*\**P* < 0.001; \*\*\*\**P* < 0.0001 one-tailed t-test. The complete list of *P* values is provided in **Table S4**.

To assess the chronic biological response to our PINE probe, we performed immunochemical staining on cross-sections of plant tissue containing implanted probes (**Materials and Methods**), so as to directly visualize lignin and suberin (two biopolymers plants produce to seal injuries). These two biopolymers accumulate robustly after damage, and can serve as reliable markers of host defense. We mapped their spatial distribution at the implantation site and quantified tissue damage: higher deposition indicates a stronger immune reaction. Biocompatibility was evaluated by comparing responses to the PINE probe versus bare metal wire. **Figure 5E** shows that after 15 days, wound-induced tissue accumulation—and lignin fluorescence — around the metal wire far exceeded that around the PINE probe. Furthermore, **Figure 5F-G** reveals the fluorescence full-width-at-half-maximum (FWHM) is ∼80 µm × 5 µm, much smaller than typical tomato stem cells (∼100–200 µm in diameter).

We assessed how implanted PINE probes affect wound healing in tomato stems by quantifying lignin levels via confocal-based histology, and analyzed five biological replicates (*n* = 5) at 3, 7-, 10-, 15-, and 20-days post-implantation. Samples contained either PINE probes or metal wires (**Figure 5H**; **Figures S4-S7**). Lignin levels remained low for the first week, rose gradually, peaked at day 10, and then stabilized—indicating progressive tissue adaptation to the probe. This time-dependent response aligns with the plant’s natural defense mechanism. Physiologically, implantation caused visible wounds within the first 7 days—consistent with mechanical injury—but by day 15, wounds were fully healed with no residual damage. Thus, the tungsten wire–assisted implantation minimizes trauma and accelerates healing, outperforming alternative methods.

We quantitatively and comparatively analyzed time-dependent biological responses after implanting PINE and metal wires. This analysis used cross-sectional imaging data from plant tissues collected at various time points post-implantation, focusing on the spatial extent (area) and fluorescence intensity of lignin at the implantation sites. Over time, lignin levels at PINE and metal wire implantation sites converged, indicating comparable tissue maturation. However, metal wires consistently produced larger wound areas than PINE (**Figure 5I, J**). Crucially, PINE can be precisely delivered into plants, enabling reliable, direct ion measurement. Thus, the data provide quantitative evidence that PINE probes are significantly less disruptive to plant tissue than conventional methods.

## 3. Discussion

We developed ultrathin, flexible PINE arrays that chronically monitor plant ion fluxes *in vivo* with high spatial and temporal resolution—overcoming key limitations of conventional techniques (bulk biochemical assays, rigid microelectrodes, and low-sensitivity fluorescence probes). Their cell-scale dimensions and mechanical compliance minimize mechanical stress and preserve cellular integrity. Biocompatible surface modifications further reduce immune-like responses in plant tissue and enhance long-term stability. Moreover, PINE probes are assembled and implanted using a method that minimizes plant damage, enabling precise, controlled placement in target regions for reliable, non-disruptive ion monitoring. Their nanoscale design offers exceptional sensitivity, selectivity, and biocompatibility—properties that advance plant physiology research, particularly in drought tolerance and symbiotic efficiency—and is requiring for the future development of plant ionomics.[44].

PINE probes with sub-cellular dimensions minimize signal dilution, allowing reliable detection of sub-micromolar ion concentrations. They will resolve transient pH-driven H⁺ fluctuations in guard cells during stomatal closure with 100-ms temporal resolution. Unlike conventional patch-clamp electrodes or rigid MEAs, PINE arrays feature a high surface-to-volume ratio—enabling rapid ion diffusion, suppressing charging artifacts, and delivering superior signal-to-noise ratios. Crucially, its ultra-thin polyimide substrate imposes minimal mechanical stress on plants during growth, preserving cellular integrity. Moreover, PINE maintains stable function during the life cycle of soil-grown tomatoes—outperforming all previous reports—and thus greatly advances studies of plant ion regulation mechanisms[47]. As a result, new applications such as tracking multiple ion fluxes become feasible. Clarifying nutrient uptake, stress responses, and symbiotic interactions allows ion dynamics to be naturally integrated into ecophysiological models—improving predictions of crop resilience to drought and salinity. Crucially, understanding ion dynamics will enable rational design of ion channels and transporters to enhance nutrient use efficiency [48].

Although PINE offers valuable tools for plant ionomics, its adoption is limited by key drawbacks. Current PINEs use ionophores for ion selectivity, yet performance in plant systems is suboptimal—for example, CNT-based ionophore-functionalized electrodes exhibit K⁺/Na⁺ cross-reactivity, reducing accuracy in complex plant tissues.[49]. In addition, many ion-selective membranes (ISMs) degrade under the extreme pH conditions common in plants (e.g., acidic vacuoles or alkaline apoplast), causing signal drift[50]. Plant tissue heterogeneity—such as vascular bundles and epidermal layers—complicates probe implantation; current PINE probes cannot penetrate dense tissues like sclerenchyma, restricting measurements to superficial or soft cells[4]. Furthermore, the lack of a cohesive network among disparate nanoelectrode arrays makes it arduous to precisely map 3D ion gradients in organs such as leaves or roots.

PINE’s future architecture will enable customizable, high-resolution electrode networks that conform to plants’ heterogeneous morphology and integrate multi-scale data—from ion channels at the cell membrane to tissue-level interactions—delivering a unified view of ion regulation. Technologically, it aims to develop more biocompatible, responsive nanomaterials. Combining advanced nanofabrication with appropriate techniques will directly address key challenges, clarify ion–biological interactions, and significantly advance plant electrophysiology. Critically, aligning technological innovation with plant biology’s inherent complexity will revolutionize ion dynamics research—and accelerate progress in sustainable agriculture and environmental resilience[3, 15, 44].

## 4. Materials and Methods

### Plant materials and growth conditions

The tomato seeds were obtained from Han Xiao. *S. lycopersicum* (cv. MicroTom) plants were grown in plastic pots containing a 1:1:1 mixture of soil, perlite, and vermiculite, under a 16-hour light/8-hour dark photoperiod at 18-22 °C. The plants were watered every two days in the phytotron facility at the CAS Centre for Excellence in Molecular Plant Sciences.

### PINE fabrication

A 6-inch fused silica wafer was used as the insulating substrate. A nickel layer was deposited as a sacrificial and release layer. A polyimide (PI) layer was then spin-coated and heat-treated for thermal stability. Chromium/gold (Cr/Au) was deposited as the metal wiring. A Cr/Ni/Au stack was patterned by photolithography to form a bonding pad for welding to the backend printed circuit board (PCB). An upper PI layer was applied, and an etching mask was defined photolithographically. Through-holes were etched based on the mask, and Cr/Au was deposited in the designated areas using photolithography and metal deposition. The flexible electrode tip was then released and encapsulated. A solid-state contact layer was electrochemically deposited on the electrode surface, followed by a potassium ion-selective membrane that allowed selective passage of K⁺ ions, completing the PINE sensor. Finally, a custom PCB was mounted on the contact pads, and the implantable section was immersed in a nickel etchant for 10 minutes to release the flexible part, yielding the final PINE array.

### PEDOT-PSS-CNT electroplating

To modify the electrode, conductive polymers doped with carboxylated multi-walled CNTs were used. The flexible array was first ultrasonically cleaned for 10 minutes. Electrode modification was performed using an electrochemical workstation (CHI 660e, Shanghai). The electroplating solution was prepared by mixing 0.1 M PSS (Sigma-Aldrich, USA), 0.01 M EDOT (Sigma-Aldrich, USA), and CNTs (XFNANO, China). A homogeneous electroplating solution was prepared by dispersing 1 mg/ml of CNTs in 0.1 M PSS and 0.01 M EDOT, followed by 1 hour of sonication to ensure uniform dispersion. Using a three-electrode setup, the modified electrode sites acted as the working electrode, with a platinum wire as the counter electrode and Ag/AgCl as the reference. Electrodeposition was carried out via chronoamperometry at 0.9 V for 100 seconds, producing PEDOT-PSS-CNT-modified electrode sites. Under the same conditions, electrode sites were also modified using only 0.1 M PSS and 0.01 M EDOT, resulting in PEDOT-PSS-modified electrodes. The original, unmodified electrode was labeled as Pristine.

### Ion-selective membrane preparation

The membrane composition consisted of 32.7 wt% polyvinyl chloride, 66 wt% 2-Nitrophenyl octyl ether, 0.3 wt% potassium tetrakis [3,5-bis(trifluoromethyl)phenyl] borate, and 1 wt% valinomycin dissolved in 2 ml of THF (total weight: 200 mg). The mixture was sonicated for 30 minutes to ensure homogeneity. Subsequently, 2 μl of the freshly prepared solution was drawn using a glass microelectrode syringe pump (RWD R480, Shenzhen, China) and deposited onto the surface of the PEDOT-PSS-CNT layer. The device was then left to dry at room temperature for 24 hours. Following drying, it was activated in 0.01 M KCl for 12 hours and subsequently stored in the same solution.

### Assembly and delivery of PINE probes

Electrochemical etching was performed in a 1 M KOH solution to sharpen 75 μm diameter tungsten microwires (W1, Shanghai Hengmi Metal Materials Co., Ltd.). The wires were cut into ∼3 cm lengths and attached to copper strips using silver epoxy resin. A voltage of 1.5 V was applied relative to a graphite counter electrode (GR001CC, GraphiteStore) to control the etching rate, with the wire tip positioned at the solution-air interface. A sharp tip typically formed within 5 minutes. The released PINE device was then connected to a sharpened tungsten wire shuttle using a bio-dissolvable adhesive (2.5% PEG 400k, Sinopharm Chemical Reagent Co., Ltd.), and the PINE array was assembled. The device was mounted on a stereotaxic instrument and implanted into the tomato stem using the tungsten wire, with insertion depth adjusted gradually. Deionized water was applied to dissolve the PEG and detach the wire, which was then removed. Dental cement was used to secure the PINE device in place on the stem.

### Simulations of plant cell micromotions

A 3D finite element model was developed to study the mechanical interactions between implanted probes and plant cells during micromotions. The plant cell was modeled as 80 μm long and wide, and 200 μm tall, with a uniform 4 μm cell wall thickness. The PI electrode measured 200 μm in length, 50 μm in width, and 1 μm in thickness, while the stainless-steel electrode was cylindrical, with a 50 μm base diameter and 200 μm length. Both were implanted 100 μm into the cell. The cell sap was modeled as an incompressible neo-Hookean material with a shear modulus of 1 kPa and a density of 1000 kg/m³. It was discretized using C3D8RH elements in ABAQUS 2025, while other components used C3D8R elements based on their mechanical properties (see **Table S2**). Before micromotion loading, the top of the probe was fixed, and the other end was left free. Lateral displacements of 0.2 μm were applied to five faces of the cell, excluding the insertion site. The probe-tissue interaction was modeled with a friction coefficient of 0.01 for tangential behavior and hard contact for normal behavior.

### Characterization of electrical properties

#### Electrochemical characterization

The electrochemical performance was evaluated using a CHI660 electrochemical workstation (Chenhua, Shanghai, China). Cyclic voltammetry (CV) was performed on Pristine, PEDOT-PSS, and PEDOT-PSS-CNT in 1 M KCl, with 10 mM [Fe(CN)₆]³⁻ added for the latter two, within a voltage range of 0 – 0.5 V. A three-electrode setup was used for *in vitro* tests: PINE as the working electrode, a platinum wire as the counter electrode, and Ag/AgCl as the reference electrode. Chronopotentiometry (CP) and water layer tests assessed the ion-selective membrane’s potential stability. For CP, samples were conditioned in 0.1 M KCl for 24 h, then ±1 nA current was applied for 100 s each while recording the potential response in 0.1 M KCl. In the water layer test, electromotive force (EMF) was measured to evaluate K⁺ response and stability. *In vivo* electrochemical tests used a two-electrode system with PINE as the working electrode and Ag/AgCl as the counter electrode.

#### EIS measurements

*In vitro* impedance testing was conducted using a standard three-electrode electrochemical system. The working electrode was a PINE, providing a stable and inert surface for reactions. A platinum wire served as the counter electrode, completing the circuit without interfering with the process. The reference electrode was Ag/AgCl, chosen for its stability and reliability. The electrolyte used was 0.1 M KCl, a common supporting electrolyte that ensures sufficient ionic strength while minimizing side reactions. EIS measurements were recorded between 1 Hz and 100 kHz to evaluate interfacial impedance across various AC conditions. This range captures low-frequency responses related to diffusion and charge accumulation, and high-frequency responses reflecting electrolyte resistance and double-layer capacitance.

#### FIB and SEM imaging

A Quorum SC7620 sputtering coater was employed to deposit a gold layer for 45 seconds at a current of 10 mA. SEM imaging was conducted using a ZEISS Gemini SEM operating at 3 kV. Prior to FIB milling performed with an FEI Scios 2 HiVac system, a protective platinum layer approximately 300 nm thick was deposited via sputtering to minimize ion beam damage to the region of interest. To examine the cross-sectional structure of the PINE, SEM images were acquired using a secondary electron detector at 2 kV, which provided clear contrast and distinct layer differentiation.

#### AFM measurement

The surface topography of PEDOT-PSS-CNT was analyzed using a Bruker Dimension Icon atomic force microscope in tapping mode under ambient conditions. The AFM uses a laser beam deflection system and closed-loop XYZ stages to measure surface roughness by detecting vertical movements of a sharp probe tip (radius <10 nm) as it scans across the sample. The scan area was 3 × 3 μm², with a resolution of 256 × 256 pixels, providing a lateral resolution of less than 1 nm per pixel, and a scan speed of 0.8 Hz. Surface roughness parameters *R_a_* (arithmetic mean deviation) and *R_q_* (r.m.s.roughness) were calculated using Gwyddion or Bruker NanoScope Analysis software.

#### Ellipsometry measurement

The thickness of the potassium ion-selective film was measured using ellipsometry (Horiba UVISEL PLUS, France) over a spectral range of 200–2000 nm. The system used a 150 W xenon lamp and a monochromator for wavelength selection. A rotating compensator at 100 kHz modulated the polarization, which was analyzed by a linear polarizer and photodetector array. Data were collected every 5 nm at a 70° angle of incidence. The *Ψ* and *Δ* spectra were analyzed using WVASE32 software with optical models: Cauchy (transparent films), Tauc-Lorentz (semiconductors), and Bruggeman (composites). Parameters were fitted to minimize RMS errors (<2° for *Ψ*, <0.5° for *Δ*). Instrumental drift was corrected using a reference SiO₂/Si wafer (*n* = 1.46, *k* = 0 at 300 K). Background signals were removed by subtracting substrate-only measurements, and polarization was calibrated using a perfect conductor mirror (*Ψ* = 0°, *Δ* = 90°).

### Histology sample preparation

To assess the effects of PINE implantation on plants, a PINE probe and a metal wire were implanted into the stems of four-week-old tomato plants. Stems were collected at 3, 7-, 10-, 15-, and 20-days post-implantation and cleaned for histochemical staining. To examine lignin and suberin accumulation at the wound site, mature stems were embedded in 5% agar and cross-sectioned using a vibratome (VT1000s, Leica). Sections were stained with 0.2% Basic Fuchsin (215597, Sigma) for 2 hours, rinsed three times with ClearSee solution, then stained with 0.01% Fluorol Yellow 088 (sc-215052, Santa Cruz Biotechnology) for 30 minutes, and washed overnight in ClearSee solution. After dual staining, sections were incubated in 0.1% Calcofluor White solution (18909, Sigma) for 30 minutes and then placed in ClearSee solution for 2 hours.

### Confocal microscopy

The stem tissue sections were mounted on glass slides and examined using a Leica SR5 confocal laser scanning microscope to visualize cell walls, lignin, and suberin. Cellulose was excited at 405 nm, with fluorescence detected in the range of 450–500 nm. Lignin was excited at 561 nm, and its fluorescence was captured between 600–650 nm. Suberin was excited at 488 nm, with emission detected between 500–550 nm. A 63×/1.4 NA oil-immersion objective was used for high-resolution imaging, while a 10×/0.40 DRY objective was employed for preview imaging. Fluorescence signals were detected using Power HyD S Detectors. The pixel size was set to 0.76 μm, with a zoom magnification of 1.5× for detailed imaging, and a scanning speed of 200 Hz. A pinhole size of 1 Airy Unit (corresponding to an axial resolution of 53.1 μm) was applied. Multichannel images were merged using LAS X software.

### Quantification of the histology

To quantify the extent of tissue damage at the implantation site, the wound area and lignin accumulation were measured as key indicators. The wound boundary in each section was manually outlined and the area calculated using a custom analysis script developed in ImageJ. The degree of lignin accumulation was indicated by pink fluorescence intensity; therefore, the corresponding fluorescence channel was isolated, normalized to eliminate background noise, and evaluated based on relative fluorescence intensity values derived from gray value measurements. Bar graphs were generated and statistically analyzed using GraphPad Prism v.8.0.1. Differences with *P*<0.05 were considered statistically significant. Filled dots represent individual data points. In all cases, the number of biological replicates is indicated as *n*. Individual *P* values for all statistical analyses are listed in **Table S4**.

### Image data analysis

Image analysis of all horizontal slices was performed using ImageJ software in combination with a custom Python script. For each image, the shortest distance from every pixel to the boundary of the PINE was calculated. Pixel intensities were then grouped into bins at 35 μm intervals based on these distances. Within each bin, the mean fluorescence intensity was calculated and subsequently normalized to a baseline, defined as the average intensity of pixels located 511 – 600 μm away from the PINE boundary. Normalized intensity profiles were averaged across samples, and the standard deviation was calculated to quantify inter-sample variability.

## Statistics

All the statistical analyses were conducted in Python (v3.13.7). One-sided Welch’s *t*-tests were employed to assess the minimized disruption caused by the PINE probe in **Figure 5I, J**. The plant sample size was selected in accordance with species-specific guidelines to ensure experimental repeatability while minimizing the use of research plants. Chronic *in vivo* tracking of electrochemical impedance spectra and electromotive force data has been reproduced across five effective sites to evaluate the electrochemical stability of the PINE probe. No plant species were excluded. PINE electrodes with impedance >2 MΩ (at 1 kHz) were deemed defective and excluded from statistical analysis. Histological analyses were performed on five randomly selected regions of each tomato stem and repeated across five independent samples at multiple time points post-implantation. Differences between PINE- and metal wire–implanted tissue sections were assessed using the ’statsmodels’ toolbox.

## Acknowledgements

We thank Prof. H. Xu, Ms. J. He, and Dr. J. Hao for their insightful input during the pilot phase. We also thank the Biochemical Sensitive Materials and Effects Research Platform at the Shanghai Institute of Microsystem and Information Technology for timely facility support during technical challenges.

## Data Availability Statement

All data are available in the main text or the supplementary information.

Python3.13.7 is available at: https://www.python.org/downloads/?pStoreID=massmutual\\n, with ‘statsmodels’ available at: https://www.statsmodels.org/stable/index.html.

Image J 1.54p is available at: https://imagej.net/ij/download/

Remaining analysis was done in customized Python script and is available upon reasonable request.

