## Supporting Information for "Ultraflexible Ion-Specific Nanoelectrodes for Long-Term Potassium Monitoring in Plants"

**This PDF file includes:**

Supporting text

Figures S1 to S7

Tables S1 to S4

Legends for Movies S1 to S2

SI References

**Other supporting materials for this manuscript include the following:**

Movies S1 to S2

### Supporting Information Text

Implantable electrodes for monitoring chemical signals in plants must be carefully designed to ensure long-term stability and accurate data collection. Two key mechanical properties are footprint—the physical area of contact with plant tissue—and bending stiffness, which determines how well the electrode conforms to or supports the surrounding biological structures. These factors are critical in determining the electrode's biocompatibility, signal fidelity, and longevity within the plant system. For instance, the footprint of an implantable electrode directly affects how it interacts with the plant's internal environment. A larger size increases the surface area for electrochemical reactions, potentially enhancing signal detection sensitivity. However, it can also cause greater mechanical disruption to plant tissue during insertion and over time, triggering stress responses such as cell death, oxidative bursts, or the deposition of lignin and suberin around the electrode. These biological reactions can insulate the electrode, reducing its ability to accurately track chemical signals like ions, hormones, or metabolites. On the other hand, a smaller electrode size causes less damage and promote better integration with the plant. However, it may also limit the electrode's capacity to capture robust and representative chemical signals, especially in heterogeneous tissue environments.

Bending stiffness determines how electrodes respond to mechanical stresses within plants. Plant tissues are dynamic, often soft and flexible, constantly growing and responding to external stimuli like wind or touch. Electrodes with high bending stiffness are rigid and less adaptable to these natural movements, thus causing mechanical strain at the interface between the electrode and the plant tissue. This strain can lead to localized damage, triggering a wound response that includes the deposition of lignin and suberin — rigid, hydrophobic polymers. Over time, this reaction can encapsulate the electrode, thus altering its ability to detect chemical signals. In contrast, electrodes with lower bending stiffness are more compliant and better adapt to natural plant tissue movements and deformations. This mechanical compatibility reduces tissue response and maintains a stable interface between the electrode and the plant. As a result, the electrode can function reliably for longer periods, allowing for consistent signal readings. Lower stiffness also improves long-term signal quality by reducing motion-related interference, which can otherwise create unstable and noisy signals.

**Calculation of bending stiffness.** To benchmark our PINE electrode, we reviewed previous studies on plant–sensor interfaces that utilized substrates with elastic moduli ranging from ultra-soft cryogels (~10–40 kPa) to PET films (~2.5 GPa), as well as rigid metallic or carbon probes (~70–230 GPa). The bending stiffness of plate-like electrodes reported in the literature was normalized as

$$D = \frac{Ewt^3}{12} \quad \textbf{(S1)}$$

where $E$ is the Young's modulus, $w$ the electrode width and $t$ its total thickness.

Cylindrical probes were processed using the following formula:

$$D = \frac{E\pi d^4}{64} \quad \textbf{(S2)}$$

where $d$ is the outer diameter, allowing for direct mechanical comparison among Polyimide films, rigid metal, silica fibers and carbon wires.

**Calculation of capacitance.** The capacitance (C) was calculated based on the cyclic voltammograms, employing the following formula:

$$C = \frac{1}{2vSV}\int_{V_i}^{V_f} I(V)\,dV \quad \textbf{(S3)}$$

where $v$ is the scan rate ($Vs^{-1}$), $S$ the total surface area of working electrode ($mm^2$), $\Delta V$ the applied voltage window (V), $V_f$ the final voltage, $V_i$ the initial voltage, and *I(V)* the voltametric current (A).

**Calculation of electroactive surface area.** The electroactive surface area was determined using the Randles-Sevcik equation：

$$I_{pa} = (2.95 \times 10^5)n^{3/2}ACD^{1/2}v^{1/2} \quad \textbf{(S4)}$$

where $I_{pa}$ (A) is the anode peak current, $n$ the number of electrons participating in the redox event, $A$ ($mm^2$) the electrode surface area, $D$ ($mm^2/s$) the diffusion coefficient, C ($mol/mm^3$) the concentration of $K_3[Fe(CN)_6]$, and $v$ (V/s) the scan rate. The diffusion coefficient of the solution containing 0.001 M of $K_3[Fe(CN)_6]$ in a 1 M KCl solution was calculated to be $D$ = $7.6 \times 10^{-4}$ $mm^2/s$.

### Figures

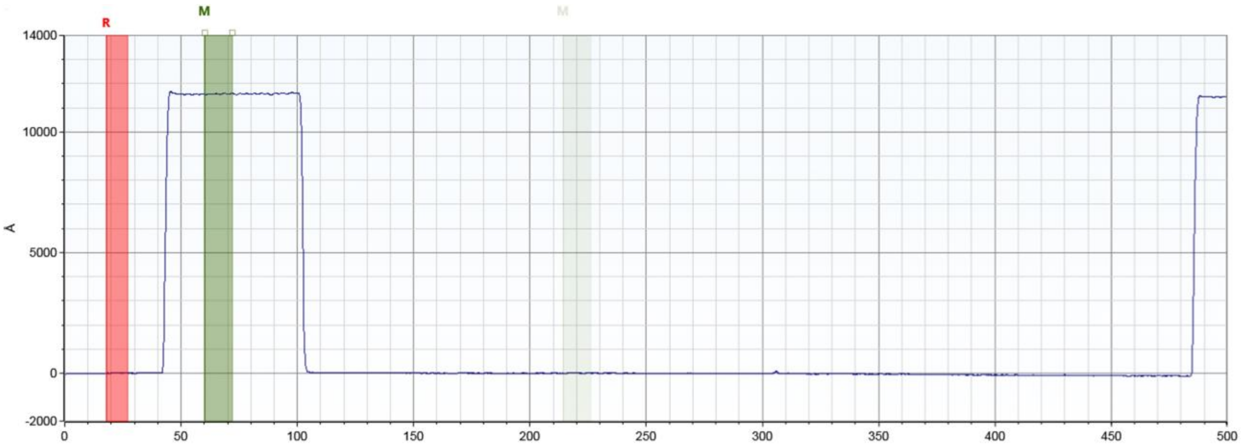


**Figure S1. Profile of the PINE probe showing an overall thickness of ~1.2 µm.**

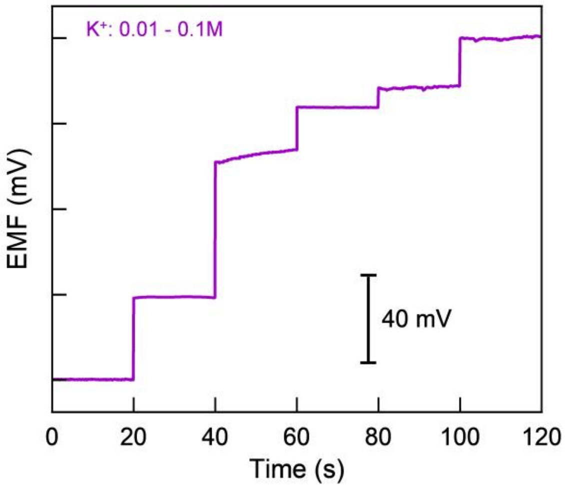


**Figure S2. Electrochemical characterization of the potentiometric response of the PINE *in vitro*.** The EMF was recorded at 20-second intervals as the concentration of KCl solution was progressively increased from 0.01 to 0.1 M.

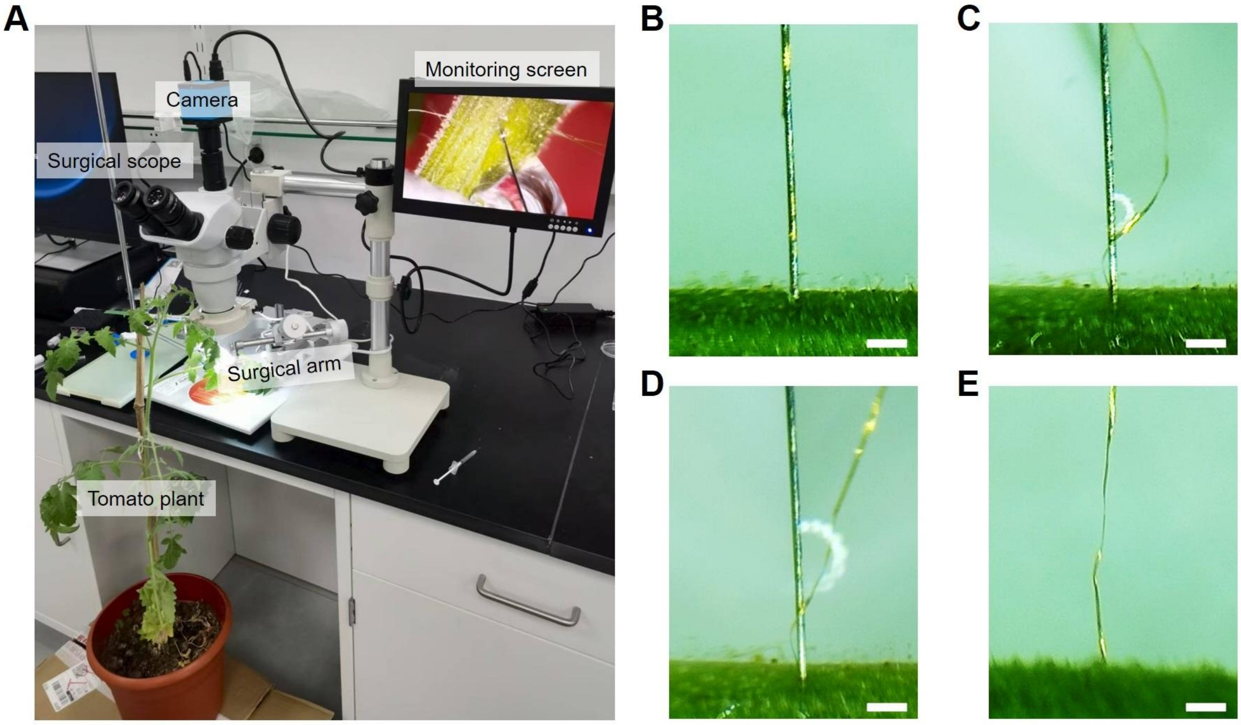


**Figure S3. Digital pictures and sequential micrographs illustrating the implantation procedures for delivering PINE into the tomato stem.** **(A)** An overview photograph of the complete setup used for implanting the PINE device. **(B)** Insertion of an assembled PINE device. **(C)** Dissolving the bio adhesive by adding DI water. **(D)** Flashing and detaching the PINE probe from the shuttle wire. **(E)** Retracting the shuttle wire from the stem. Scale bars, 500 μm.

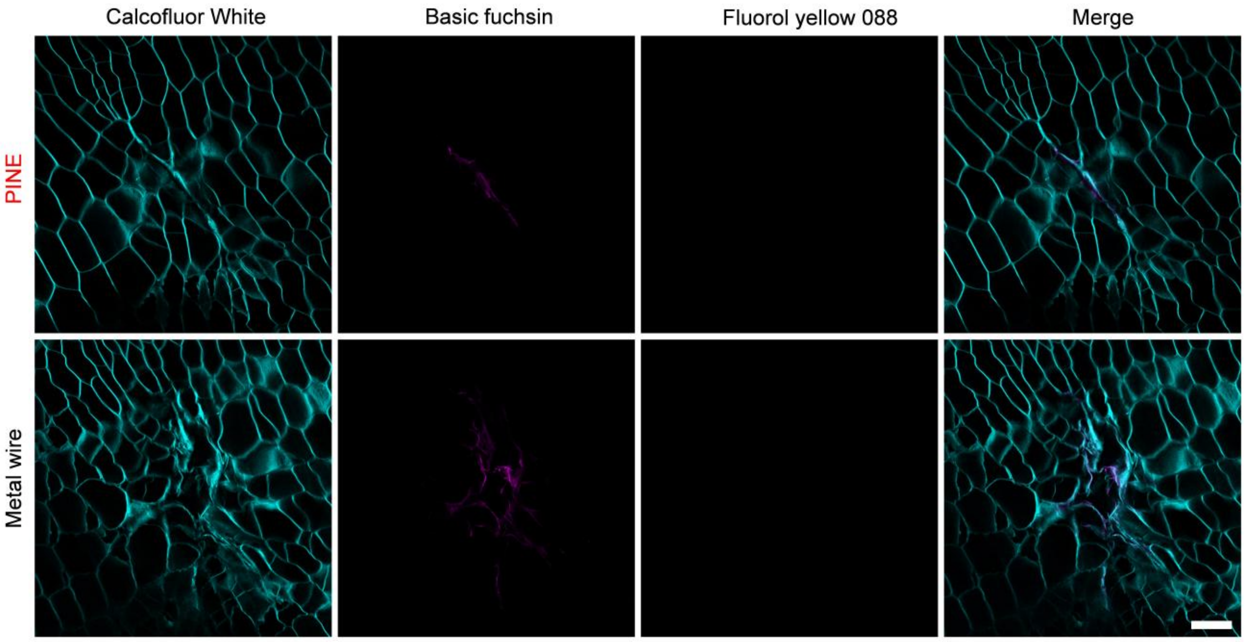


**Figure S4. Histology of PINE/plant stem cell interface.** Confocal micrographs depicting cell wall, suberin, and lignin staining, along with merged channel images, from cross-sectional slices of a PINE probe (top) and the surrounding area of a 75-μm-diameter steel electrode (bottom), both implanted in the tomato plant stem 3 days post-implantation. Scale bar, 100 μm.

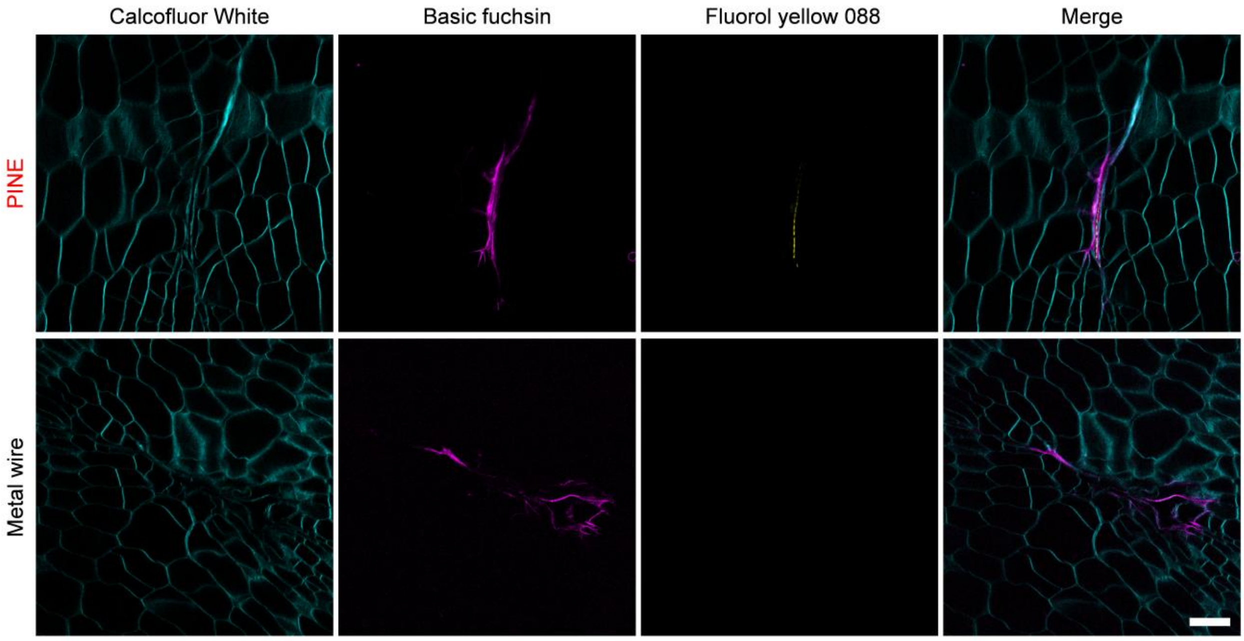


**Figure S5. Histology of PINE/plant stem cell interface.** Confocal micrographs depicting cell wall, suberin, and lignin staining, along with merged channel images, from cross-sectional slices of a PINE probe (top) and the surrounding area of a 75-μm-diameter steel electrode (bottom), both implanted in the tomato plant stem 7 days post-implantation. Scale bar, 100 μm.

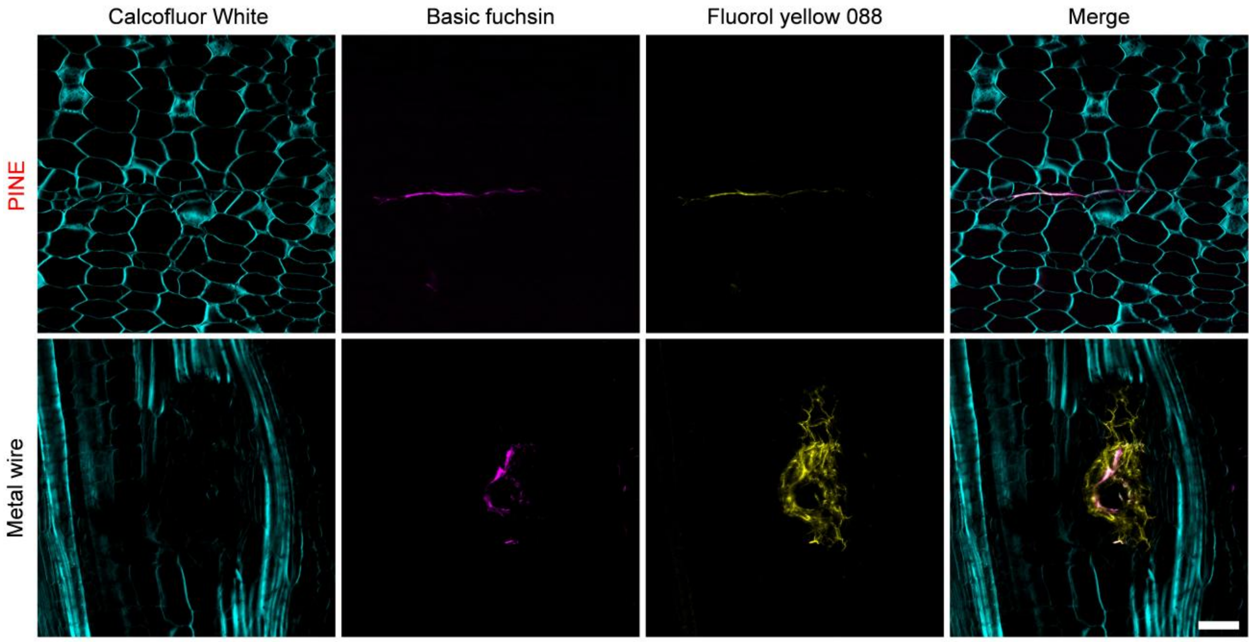


**Figure S6. Histology of PINE/plant stem cell interface.** Confocal micrographs depicting cell wall, suberin, and lignin staining, along with merged channel images, from cross-sectional slices of a PINE probe (top) and the surrounding area of a 75-µm-diameter steel electrode (bottom), both implanted in the tomato plant stem 10 days post-implantation. Scale bar, 100 µm.

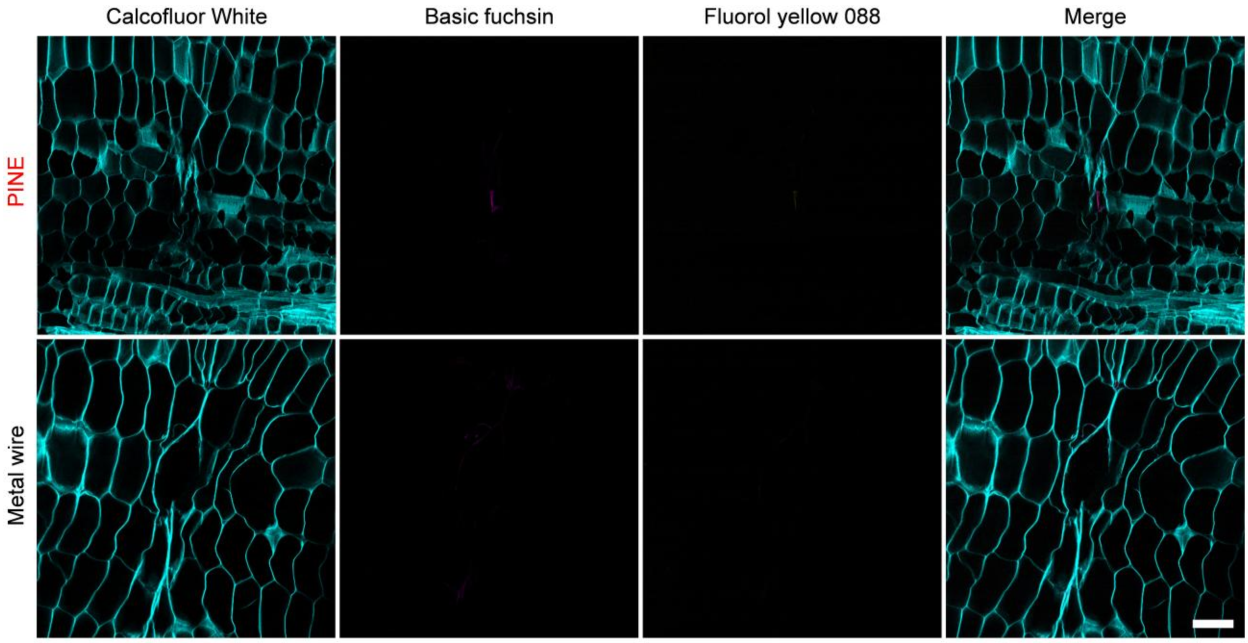


**Figure S7. Histology of PINE/plant stem cell interface.** Confocal micrographs depicting cell wall, suberin, and lignin staining, along with merged channel images, from cross-sectional slices of a PINE probe (top) and the surrounding area of a 75-µm-diameter steel electrode (bottom), both implanted in the tomato plant stem 20 days post-implantation. Scale bar, 100 µm.

### Tables

**Table S1. Types and dimensions of state-of-the-art electrodes utilized for *in vivo* chemical sensing in plants.**

| *Probe type* | *Probe materials* | *Chemicals* | *Implantation site* | *Cross sectional area ($mm^2$)* | *Shape & Avg. size (mm)* | *Ref.* |
|---|---|---|---|---|---|---|
| **Rigid** | Steel | $Na^+$ | Tomato stem | 0.011 | Cone, tip dia. ~$2.3\times10^{-3}$, medial dia. ~0.12 | [1] |
| **OECT** | Cryogel | $K^+$ | Tomato stem | 6.25 | Square 2.5 × 2.5 | [2] |
| **Rigid** | Ti wire | abscisic acid | Cucumber fruits, juices, grapes, radishes, Arabidopsis leaf | 0.28 | Cylinder, Dia. ~0.6 | [3] |
| **Rigid** | Graphite rod | L-tryptophan | Tomato and cherry tomato fruits | 3.14 | Cylinder Dia. ~2 | [4] |
| **Flexible** | PET | Sugar | Tomato fruits | 3.03 | 10.1 × 0.35 | [5] |
| **Flexible** | Resin | pH | Leaf | $0.7\times10^{-3}$ | Micro needle Dia. ~0.03 | [6] |
| **OECT** | Tentile absorbent or polumer microfibers | Glucose | Tomato stem | 0.79 | Cylinder Dia. ~1 | [7] |
| **Rigid** | Carbon fiber | Ascorbic acid | Lemon, cactus | $0.85\times10^{-3}$ | Cylinder Dia. ~0.033 | [8] |
| **OECT** | Cotton | NaCl | Cactus | 1.77 | Cylinder Dia. ~1.5 | [9] |
| **Rigid** | Silica fiber | ABA, IAA, GA3 | Aloes | 0.43 | Cylinder Dia. ~0.74 | [10] |
| **Flexible** | PI | $K^+$ | Tomato stem | $4\times10^{-5}$ | 1μm × 40μm | PINE |

**Table S2. Mechanical property parameters of materials used in finite element analysis models.**

| *Material* | *Young's modulus* | *Poisson's ratio* |
|---|---|---|
| **PI** | 3 GPa | 0.35 |
| **Stainless steel** | 193 GPa | 0.30 |
| **Cell wall** | 2.3 GPa | 0.4 |

**Table S3.** Comparison of $K^+$ detection performance among state-of-the-art probes.

| ***Probe type*** | ***Functional materials*** | ***Substrate materials*** | ***Linear range (M)*** | ***Limits of detection (LODs)*** | ***Chronic stability*** | ***in vivo, real-time detection*** | ***Ref.*** |
|---|---|---|---|---|---|---|---|
| **OECT** | PEDOT:PSS+DMSO | photographic paper | $10^{-4}$-$10^{-1}$ | $10^{-4}$ | / | / | [11] |
| **OECT** | PEDOT:PSS | Polyethylene naphthalate (PEN) substrate | $4.92\times10^{-6}$-0.1 | $4.92\times10^{-6}$ | / | / | [12] |
| **OECT** | PEDOT:PSS | paper | $1\times10^{-6}$-$0.1\times10^{-6}$ | $0.1\times10^{-6}$ | / | / | [13] |
| **OECT** | PSS:Na | silicon/silicon dioxide ($Si/SiO_2$) substrate | $10^{-3}$-$10^{-2}$ | $10^{-3}$ | / | / | [14] |
| **OECT** | PEDOT:PSS，BBL | glass slide | $4.3\times10^{-3}$-$9.6\times10^{-3}$ | $4.3\times10^{-3}$ | / | / | [15] |
| **OECT** | PEDOT:PSS | glass wafer | $1\times10^{-3}$-0.1 | $1\times10^{-3}$ | / | / | [16] |
| **OECT** | PEDOT:PSS | thermoplastic polyurethane (TPU) | $1.25\times10^{-3}$-0.1 | $1.25\times10^{-3}$ | / | / | [17] |
| **OECT** | PEDOT:PSS | glass substrates | $10^{-6}$-$10^{-1}$ | $10^{-6}$ | / | / | [18] |
| **OECT** | PEDOT:PSS | PI | $10^{-5}$-$10^{-1}$ | $10^{-5}$ | / | / | [19] |
| **OECT** | PEDOT:PSS | cryogels | / | $0.3\times10^{-6}$ | 2 months | √ | [20] |
| **SC-ISE** | reduced graphene oxide | glassy carbon | $10^{-5.03}$-$10^{-1.14}$ | $10^{-5.03}$ | / | / | [21] |
| **SC-ISE** | / | graphene paper | $10^{-5.5}$-$10^{-1}$ | $6\times10^{-7}$ | 17 h | / | [22] |
| **SC-ISE** | PANI-DNNSA | graphene paper | $10^{-6}$-$10^{-1}$ | $2.5\times10^{-6}$ | 24 h | / | [23] |
| **SC-ISE** | PEDOT-HQ | glassy carbon | $10^{-7}$-$10^{-1}$ | $2\times10^{-7}$ | 24 h | / | [24] |
| **SC-ISE** | carbon black | SPEs(screen-printed electrode) | $10^{-5}$-$10^{-1}$ | $1\times10^{-5}$ | 4 weeks | / | [25] |
| **SC-ISE** | Ni-HAB MOF | glassy carbon | $10^{-6}$-1 | $1.9\times10^{-6}$ | 20 h | / | [26] |
| **SC-ISE** | β-CD/RGO | glassy carbon | $10^{-7}$-$10^{-1}$ | $10^{-6.2}$ | / | / | [27] |
| **SC-ISE** | β-CD/RGO | CC ink/PET | $10^{-5.35}$-$10^{-1}$ | $10^{-5.35}$ | / | / | [27] |

| | | | | | | | |
|---|---|---|---|---|---|---|---|
| **SC-ISE** | CB/Nano 19 | PCL-PS TPE /Thermoplastic electrodes (TPEs)/Nano19 | $10^{-4}$-$10^{-1}$ | $10^{-4}$ | / | / | [28] |
| **SC-ISE** | Polydioxythiophene layer | glassy carbon | $10^{-6}$-$10^{-1}$ | $10^{-6}$ | / | / | [29] |
| **SC-ISE** | Carbon nanofibers /nickel–cobalt nanoparticles /CNT | glassy carbon | $10^{-6}$-$10^{-1}$ | $10^{-6.3}$ | 15 h | / | [30] |
| **SC-ISE** | Graphene | polyethylene terephthalate (PET) | $10^{-4}$-$10^{-1}$ | $10^{-4.28}$ | 15 h | / | [31] |
| **SC-ISE** | Au nanoparticle/siloxene/graphene/poly(dimethylsiloxane) | PDMS | $10^{-5.5}$ | $8.1\times10^{-5.7}$ | / | / | [32] |
| **SC-ISE** | PEDOT-PSS-CNT | PI | $10^{-10}$-$10^{-2}$ | $10^{-8}$ | 48 days (life-long) | √ | PINE |

**Table S4. Individual *P*-values for statistical (two-tail *t*-test) presented in Figure 5.**

| *Days post-implantation* | *p-value for the area* | *Days post-implantation* | *p-value for the fluorescence intensity* |
|---|---|---|---|
| **3 Day** | $4.48 \times 10^{-4}$ | 7 Day | 0.0021 |
| **7 Day** | $1.33 \times 10^{-4}$ | 10 Day | 0.7643 |
| **10 Day** | $2.60 \times 10^{-3}$ | 15 Day | 0.0037 |
| **15 Day** | $2.53 \times 10^{-3}$ | 20 Day | 0.3963 |
| **20 Day** | $7.21 \times 10^{-5}$ | | |

**Movie S1 (separate file). Implantation of an assemble PINE device into the tomato stem**. Scale bar, 500 µm.

**Movie S2 (separate file). Confocal images of the PINE/plant interface showing an intact cellular network around the PINE contacts**. Scale bar, 100 µm.
